# High-throughput Genome Wide CRISPR Knock Out mechanical sort identifies genes driving metastatic cancer cell softening

**DOI:** 10.64898/2026.02.12.705447

**Authors:** Katherine M Young, Curtis Dobrowolski, Nicholas Stone, Kalina Paunovska, Sydney Bules, Kelly Ahkee, James Hankish, Alex Chapman, James E Dahlman, Todd A Sulchek, Cynthia A Reinhart-King

## Abstract

Cell mechanics can serve as an important biomarker for cell state and phenotype, such as metastatic ability. While some molecular mechanisms underlying cell mechanical properties have been investigated through targeted analyses, a genome-wide study of human genes and gene networks that modulate cell biophysical properties has not been attempted. In this work, we combined a microfluidic stiffness-based sorting device with a genome-scale CRISPR knockout (GeCKO) screen in order to investigate the effect of individual gene knockouts on cell stiffening and cell softening across the entire protein-coding genome. We processed approximately 150 million Cas9-expressing ovarian cancer cells that had been transduced with a library of 76,000 single guide RNAs (sgRNAs) against the 19,000 protein-coding genes in the genome. The cells were sorted into 5 mechanical subsets. We identified 7 gene knockouts that were significantly depleted in the softer subsets and over 700 gene knockouts that were significantly enriched in the stiffer subsets. Of these significant genes of interest, we selected 3 genes that were highly expressed in our ovarian cancer cell line with greater than 100-fold enrichment in the stiff outlet and resulted in significant changes in ovarian cancer patient survival. These genes, *PIK3R4, CCDC88A,* and *GSK3B*, when knocked out result in a significant and predicted increase in cell stiffness. This study is the first to explore the relation between human gene expression and cell mechanics at the genome-scale to generate datasets at the intersection between cell genotype, mechanotype, and phenotype for metastatic cancer cells. The method could also be applied to study the effect of genes on other biophysical cell processes as well as for identifying pathways for the control of cellular mechanics across many cell types.

## Introduction

The process of metastasis, which causes a majority of all cancer-related deaths, involves the collection of mutations that allow primary tumor cells to modify their adhesive and mechanical properties, undergo the epithelial-to-mesenchymal transition (EMT), migrate away from their tumor, degrade the extracellular matrix (ECM), migrate to neighboring tissues, and undergo mesenchymal-to-epithelial transition (MET) to form a new metastatic tumor. EMT causes cells to change from polarized and epithelial-like to being spindly, motile, and mesenchymal-like^1^. This transition occurs with the loss of cell-cell interactions, the activation of certain transcription factors, changes in surface protein expression, a reordering of the internal cytoskeleton, and production of enzymes that degrade the extracellular matrix^2–5^.

Many of the changes described above, including the invasion and migration of metastatic cells, are associated with the mechanical properties of cells. The processes of EMT and MET reorganize the cell’s cytoskeleton, the proteins within a cell which provide structural support and allow cell motility^4,6^. Beyond the processes of EMT/MET, cancer cells undergo other mechanical processes that affect their extracellular microenvironment in addition to the mechanics of the cell itself^7^. Cell softening has been identified in many motile cell types and it has been shown for many cancer types an increase in metastatic potential correlates with a decrease in cell stiffness^6,8–10^. In addition to the importance of mechanics in cancer cell behavior, cell mechanics play an important role in cell viability, chemoresistance, sickle cell transport, leukocyte demargination, endothelial tube formation, stem cell differentiation, liver fibrosis, and fibrogenesis^11–20^. Because of the relationship between disease state and cell mechanics, cell stiffness has been suggested as an important biomarker to measure cell migratory or invasive capacity. There is currently a gap in our understanding of the molecular and network mechanisms driving this phenomenon. Understanding how cell mechanics integrate the signals of many molecular pathways may enable its use as a more accurate biomarker as well as to provide possible targets for therapeutics.

A limited subset of genes that affect cell mechanics have been studied. These studies have been limited in throughput and scope of the number of genes that could be assessed. At the single cell level, we previously demonstrated a novel low throughput methodology combining atomic force microscopy (AFM) and targeted single cell RT-qPCR to correlate gene expression for a subset of 96 genes of interest with cell mechanical properties across ovarian cancer cells of different metastatic potential^21^. A targeted RNA interference screen limited to 214 phosphatase and kinase genes in *Drosophila* was completed using a high throughput mechanical screening microfluidic technology known as real-time fluorescence and deformability cytometry (RT-FDC)^22^. This study identified gene knockdowns that resulted in significant changes in a cell’s mean Young’s modulus, including hits that were known regulators of cell mechanics and mitosis as well as novel mechanisms of mechanical modification. Another study has used population-level data of cell mechanotype and RNA sequencing or microarray data across five biological systems to create an in silico model to predict putative cell mechanics regulators^23^. While these studies represent an important basis for studying the role of gene expression in cell mechanics, a high-throughput genome-wide study of human genes and gene networks that modulate cell biophysical properties has not yet been attempted.

The use of genome-wide CRISPR knockout (GeCKO) pooled screens has allowed researchers to start exploring the connection between a cell’s genotype and various phenotypes, including chemoresistance, gene essentiality, *in vivo* metastasis, and immune activation^24–28^. The GeCKO process delivers a pooled lentiviral library of single guide RNAs targeted to over 19,000 genes (approximately every protein-coding gene in the human genome) to a Cas9-expressing cell line. A low multiplicity of infection (MOI) is used so that statistically each cell will contain at most one CRISPR knockout^24^. Cells with the single gene knockouts are screened for a phenotype of interest and sequencing is used to compare the sgRNA distribution before and after the screen to identify which genes are related to the applied screening pressure. Screens suitable for the GeCKO process should be sensitive to the phenotype of interest and of sufficient throughput to process tens of millions of cells needed for the sequencing analysis.

To understand gene networks that control cell mechanics and their role in metastatic potential, we used a high-throughput microfluidic mechanical screen in conjunction with a genome-scale CRISPR knockout strategy to uncover the genetic and molecular mechanisms that allow cells from an inherently heterogeneous tumor population to change their mechanical properties (Figure 1). The microfluidic sorting platform we use consists of a series of diagonal ridges that cells must deform underneath as they are flowed through the device. Softer cells easily deform underneath each ridge and their trajectories through the device are unaffected by the ridge resulting in sorting to outlets below the midline of the device (labeled outlets 1 and 2). Stiffer cells resist deformation and are deflected by each ridge resulting in positive y-direction translation and sorting to outlets above the midline of the device (labeled outlets 3, 4, and 5). This device has been optimized to process tens of millions of cells^29^ and has been used to sort cells that differ in stem cell pluripotency, peripheral blood mononucleated cell identity, photoreceptor differentiation, ascitic metastatic potential, and viability ^17,30–34^. This study is the first to apply a genome-wide screen to identify modifiers of cell mechanical properties and may play a significant role in the understanding metastasis, which leads to over 8 million deaths annually worldwide. The development of this new method can be applied to uncover other molecular mechanisms of cell migration and several other processes, including stem cell differentiation, immune cell development and activation, mechanotransduction, and wound healing.

**Figure 1.**
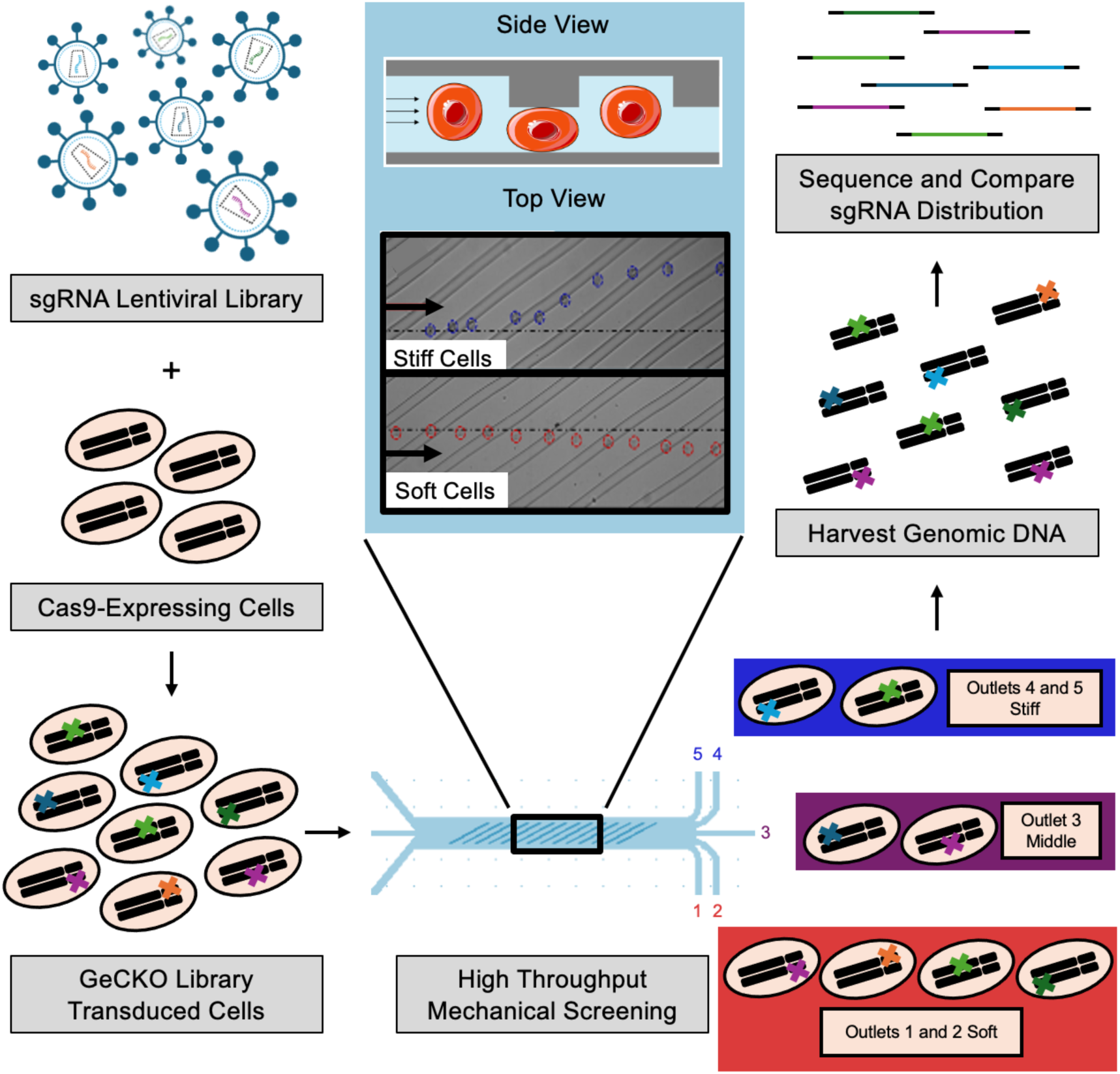
Mechanical screening to test the effect of single gene contributions to cell deformability across the whole genome. Cancer cells expressing Cas9 are transduced with a lentiviral library containing sgRNAs across the entire protein-coding genome at a low MOI to allow for single gene knockouts. Antibiotic selection ensures only cells with gene knockouts are included in mechanical screen. Next, the library of cells with single gene knockouts are processed with a microfluidic stiffness-based cell sorting device into 5 mechanical subsets. The inset shows a side and top view of the ridged region of the device where cells must deform underneath each diagonal ridge. Softer cells deform easily underneath the ridges and follow hydrodynamic streamlines to provide a slight negative deflection towards outlets 1 and 2. Stiffer cells resist deformation and are deflected along the diagonal ridge resulting in trajectories towards outlets 4 and 5. Genomic DNA is harvested from each mechanical subset, the sgRNA regions are amplified and sequenced, and the distribution of sgRNAs before and after the mechanical screen is used to determine the correlation of gene knockouts with mechanical subsets.

## Results

### Stiffness-based sorting microfluidic device can separate GeCKO transduced cells

To use a 2-vector lentiviral system for the GeCKO screen, we first established an ovarian cancer cell line that expressed Cas9. We confirmed the successful integration of a gene cassette containing Cas9 and GFP into HEY A8 cells through viral transduction by introducing a sgRNA against GFP. Using flow cytometry to compare the GFP expression level of HEY A8-Cas9-GFP cells after transfection of sgGFP, we observed a significant reduction in the number of GFP+ cells after the addition of 50, 150, and 250 ng of sgGFP (Supplemental Figure 1a). Additionally, we observed a significant reduction in the mean fluorescence intensity of the transduced cells (Supplemental Figure 1b). Together, these data confirms the successful integration of the Cas9 cassette that is able to functionally edit genes when transduced with sgRNA. As final confirmation, we sequenced the GFP region of the cells after gene editing and observed a 25-60% total insertion-deletion efficiency at the cut site as determined using TIDE analysis (Supplemental Figure 1c). The introduction of the Cas9-GFP cassette did not have an effect on the mechanical properties of the cell line (Supplemental Figure 1d), so all mechanical changes observed with sorting by the microfluidic device were attributed to the gene knockouts and not the addition of the Cas9-GFP cassette. Additionally, the cells maintained GFP expression and Cas9 expression after the cells were continually cultured and stored in liquid nitrogen for multiple years.

**Supplemental Figure 1.**
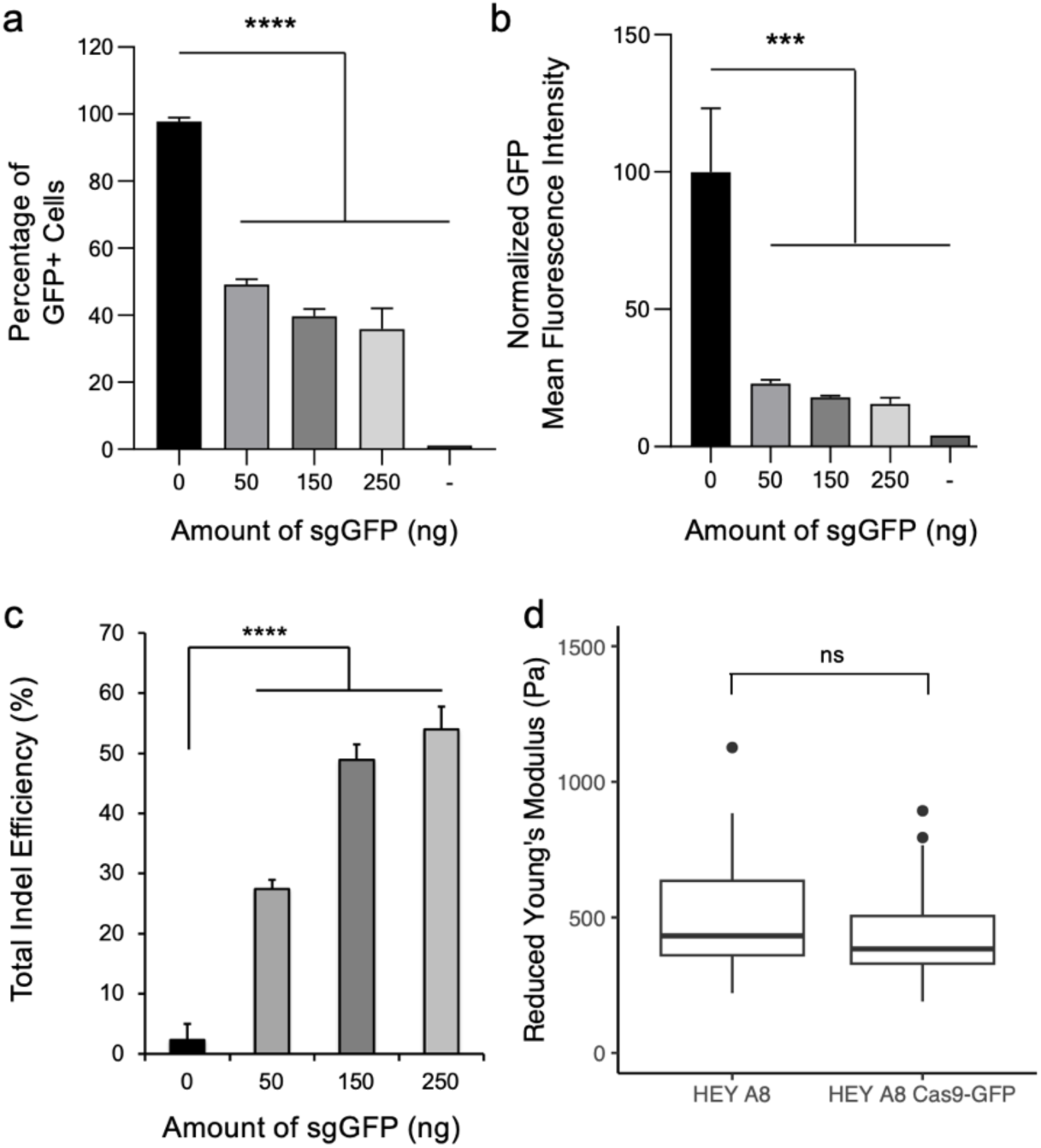
CasG-expressing cells capable of gene editing. a) The addition of sgGFP to Cas9 and GFP expressing HEY A8 cells results in the significant reduction GFP+ cells as measured by flow cytometry (Tukey HSD post-hoc test, ****, p < 0.0001). “–“ indicates cells from the HEY A8 cell line that do not express Cas9 and GFP. b) The addition of sgGFP to HEY A8-Cas9-GFP cells also results in significant reduction of mean fluorescent intensity (MFI) of each sample implying successful knockout of GFP region (Tukey HSD post-hoc test, ***, p < 0.001). “–“ indicates cells from the HEY A8 cell line that do not express Cas9 and GFP. c) TIDE analysis to determine the insertion-deletion efficiency showed significant increase after the addition of varying amounts of sgGFP to HEY A8-Cas9-GFP cells (Tukey HSD post-hoc test, ****, p < 0.0001). d) Atomic force microscopy shows introduction of the Cas9-GFP cassette did not affect the reduced Young’s modulus of the cell line. (n=40 per cell line, Welch two-sample t-test, p > 0.05.)

After establishing our Cas9 cell line, we tested the feasibility that single gene knockouts could result in detectable changes in cell mechanics and lead to sorting in the microfluidic device. High-speed video analysis was used to track cells while sorting to five cell outlets due to deflection at each ridge (Figure 2a). We transduced cells with the GeCKO lentiviral sgRNA library and took AFM measurements comparing the stiffness of cells with single gene knockouts and non-transduced Cas9-expressing cells. A significant difference in stiffness was measured between these two populations (p=0.0029, n=150 for each condition) with 5% of the GeCKO cells exhibiting measured stiffness higher than the maximum control cell stiffness (Figure **Error! Reference source not found.**2b). The 5% of GeCKO-transduced cells is estimated to represent 3,800 sgRNAs or 950 gene knockouts that could potentially increase the cell stiffness. Additionally, there was an increased variance of the single gene-knockout cell group compared to the control cell population (sd = 350.46 vs 180.21, Brown-Forsythe Test, p=0.002367), implying the GeCKO transduction process results in an increased heterogeneity of cell mechanics that could be detected by our microfluidic device.

**Figure 2.**
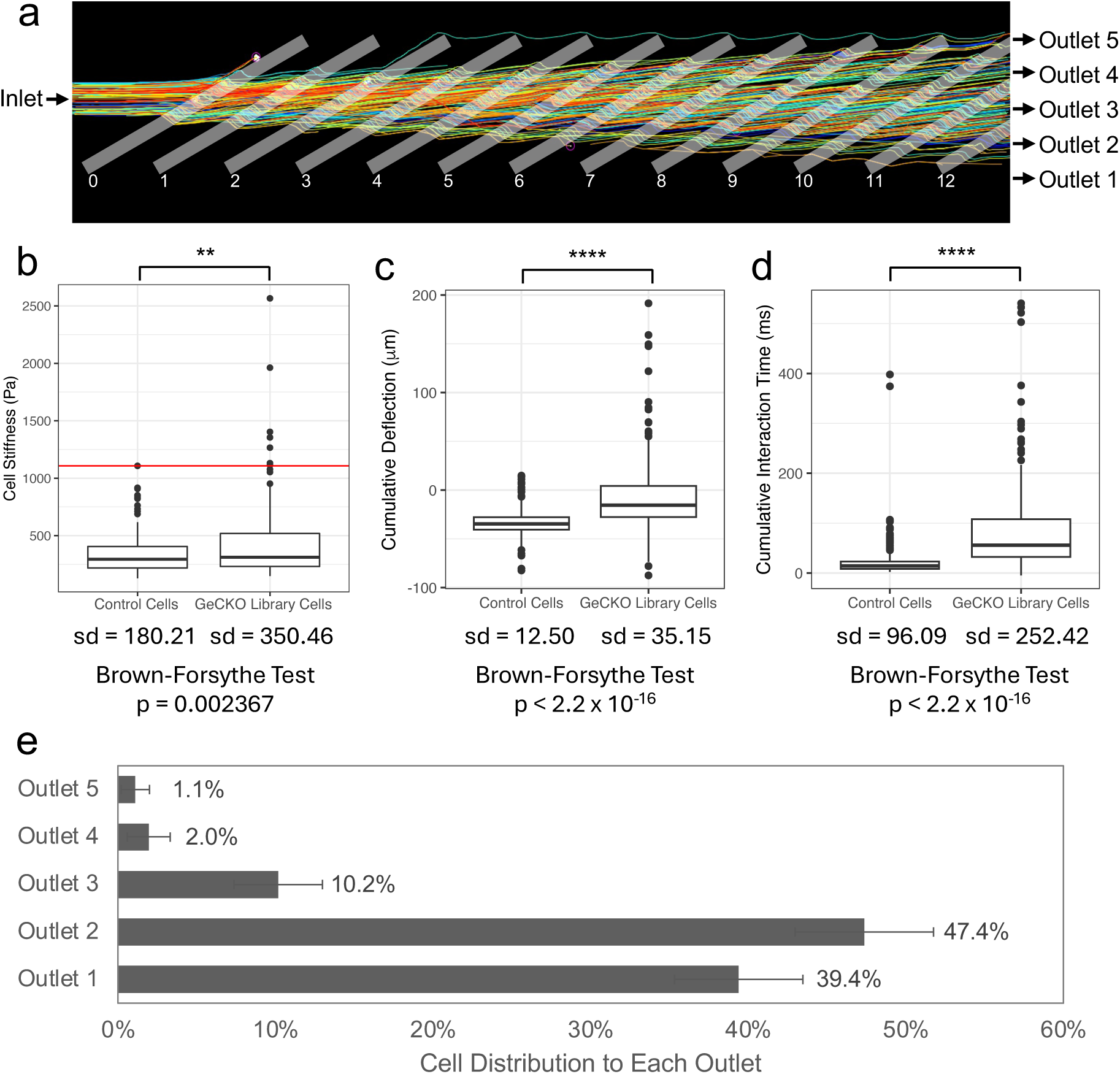
Cell mechanical changes due to single gene knockouts are sorted by microfluidic device. a) Example cell trajectories from high speed video cell tracking. Outlines were added of diagonal ridges added along with cell inlets and outlets exist in device. b) AFM measurements show GeCKO transduction leads to statistically distinct cell populations with a wider range of observed stiffnesses (**, p < 0.01, Welch two sample t-test, n = 150 per sample). The red line denotes the highest Young’s modulus value of the control cell population; 5% of the GeCKO transduced cells fell above this maximal stiffness line. The standard deviation (sd) for each group is reported along with p-value from Brown-Forsythe test comparing group variance. c) High-speed video analysis was used to track the deflection of the control and GeCKO-transduced cells in our microfluidic device from its starting to ending position after encountering each ridge. The cumulative deflection reported is the sum of deflections over the first 5 ridges. The GeCKO-transduced cells show a wider variation in cell trajectories with more cells deflected toward Outlet 5 compared to the control cells (****, p < 0.0001, Welch two sample t-test, n > 280 cells). The standard deviation (sd) for each group is reported along with p-value from Brown-Forsythe test comparing group variance. d) High-speed video analysis was used to observe the time of interaction of control and GeCKO-transduced cells in our microfluidic device with each ridge. The GeCKO cells show a wider variation in interaction time as well as increased interaction time as compared to the control cells (****, p < 0.0001, Welch two sample t-test). The standard deviation (sd) for each group is reported along with p-value from Brown-Forsythe test comparing group variance. e) Average percentage of total sorted cells that were deflected to each outlet over four bio-replicate sorts. Outlet 1 indicates the softest subset while Outlet 5 indicates the stiffest subset. Error bars denote standard deviation between the bio-replicates.

To test the behavior of these transduced cells in the microfluidic device, we used high-speed video analysis custom software to track the locations of cells deflected at each diagonal ridge. Cells with single gene knockouts showed an increased cumulative deflection (p < 2.2×10^-16^, Welch two-sample t-test), an increased variance of deflection (p < 2.2×10^-16^, Brown-Forsythe test), increased cumulative interaction time with the ridges (p < 2.2×10^-16^, Welch two-sample t-test), and an increased variance of interaction times (p < 2.2×10^-16^, Brown-Forsythe test) (Figure 2c and Figure 2d). The cumulative deflection and duration of interaction time in the device determines the trajectory of each cell to different sorting outlets. Therefore the introduction of single gene knockouts throughout the genome with the GeCKO lentiviral library resulted in changes in cellular biomechanical properties that are sortable.

Four separate GeCKO mechanical screens were conducted with 20 – 67 million transduced cells collected per experiment. In total, approximately 150 million cells were processed and collected in five mechanical subsets. The majority of the cells followed the fluid streamlines through the device to output to the soft outlets, outlet 1 and outlet 2. Approximately 3% of the transduced were deflected to the stiff outlets, outlet 4 and outlet 5 (Figure 2e). This amount is consistent with the observation that approximately 5% of cells that receive a single gene knockout will result in cell stiffening. On average, we processed cells at a rate of 6.2 million cells/hour, and the duration of processing time ranged from 4.75 hours to 11.5 hours. Sorted cell subpopulations were stored on ice until cells were pelleted and flash frozen before gDNA extraction.

### DESeq2 analysis reveals hundreds of genes enriched in mechanically sorted subpopulations

After extracting the genomic DNA from all samples, the 20 nucleotide sgRNA target region was amplified with Illumina adapter sequences and sequenced using next generation sequencing. High quality scores for the first 20 base pairs corresponds with successful sequencing of the variable sgRNA region, while a drop in quality scores after 20 base pairs corresponds to a constant adapter region with low cycle-to-cycle variability (Supplemental Figure 2a). The Gini Index was calculated to determine sgRNA distribution before and after the mechanical screen, with a low Gini Index indicating more even sgRNA distribution and a high Gini Index indicating a higher selection of unique sgRNAs. The Gini Index for the pre-screen cell inlet sample was similar to that of samples of the post-screen soft Outlets 1 and 2, while the middle Outlet 3 and stiff Outlets 4 and 5 demonstrated increased Gini indices (Supplemental Figure 2b). This implies a positive selection process in which knockouts that resulted in cell stiffening are significantly enriched in Outlets 3, 4, and 5 dominating the population. When comparing each of the bioreplicates of the GeCKO mechanical screen, the Outlet 5, Outlet 4, and Outlet 3 samples clustered distinctly by outlet, while the Outlet 2, Outlet 1, and Inlet samples clustered similarly with one another (Supplemental Figure 2c). This again highlights the positive selection of cell-stiffening gene knockouts.

**Supplemental Figure 2.**
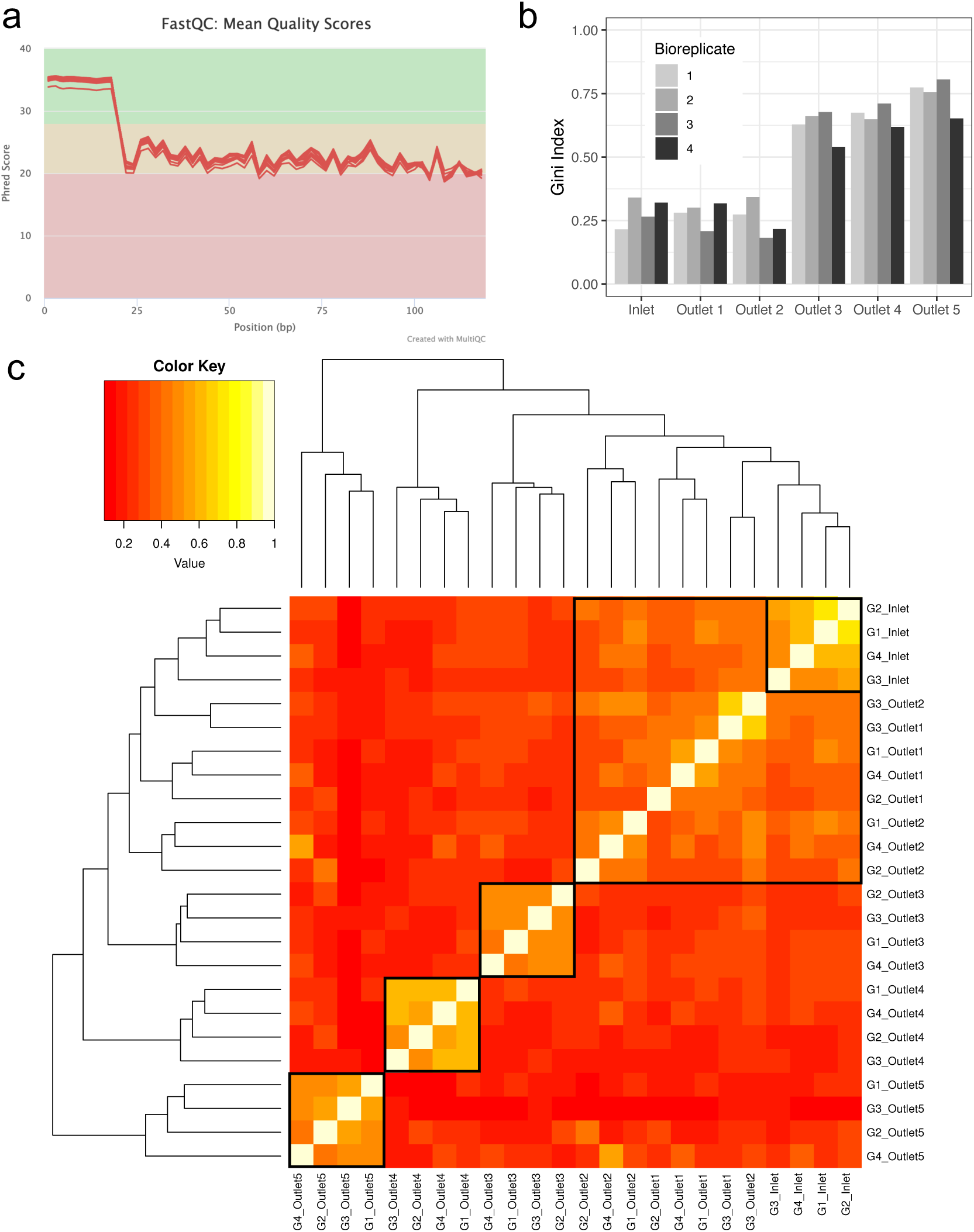
Quality control for DNA sequencing demonstrates successful amplification of sgRNA regions and positive selection for Outlets 3, 4, and 5. a) FastQC results show mean read quality Phred scores indicating high quality reads for first 20 base pairs corresponding with the 20 bp long variable sgRNA region. b) Barplot of Gini index shows increase in positive selection for Outlets 3, 4, and 5. A low Gini index indicates more even sgRNA distribution whereas a high Gini index indicates higher selection of unique sgRNAs. c) Heatmap and hierarchal clustering of samples shows clustering of biological replicates as well as distinct nature of Outlets 3, 4, and 5 compared to similarity between Outlets 1, 2, and Inlet.

Using DESeq2 to compare the pre- and post-screen distribution of each sgRNA, we identified hundreds of genes differentially distributed in each of the different mechanical subsets as compared to the inlet distributions (Figure 3a). From the softest outlet, Outlet 1, 4 gene knockouts were significantly depleted, while no gene knockouts were significantly enriched (Figure 3b). From the soft outlet, Outlet 2, 3 gene knockouts were significantly depleted and again no gene knockouts were significantly enriched (Figure 3c). From the middle outlet, Outlet 3, 111 gene knockouts were significantly enriched and 2 gene knockouts were significantly depleted (Figure 3d). From the stiff outlet, Outlet 4, 362 gene knockouts were significantly enriched and 4 gene knockouts were significantly depleted (Figure 3e). From the stiffest outlet, outlet 5, 348 gene knockouts were significantly enriched, but no gene knock outs were significantly depleted (Figure 3f). Because of the sorting efficiency of the multiplex device^29^, it is understandable that we would observe more enriched genes in Outlets 3 through 5, as we previously observed high enrichment and purity of stiffened cells. To further understand the gene knockouts that resulted in sortable changes in mechanical properties, we classified each gene into different functional categories depending on the frequency of terms that were associated with their gene ontology (GO) annotations (Supplemental Figure 3a). While there were many genes associated with the functional categories of DNA and RNA metabolism, signaling, metabolism, and transport, we were particularly interested in exploring gene knockouts related to the cytoskeleton, adhesion and migration, the cell cycle, and specific signaling pathways including the MAPK, Wnt, Notch, PI3K, JNK, and TGFb signaling pathways (Supplemental Figure 3b). We further narrowed down our list of genes of interest by focusing on genes that are in the top tercile of gene expression for the HEY A8 cell line (Supplemental Figure 3c) and that also had a sgRNA fold change of over 100-fold (Figure 3g).

**Figure 3.**
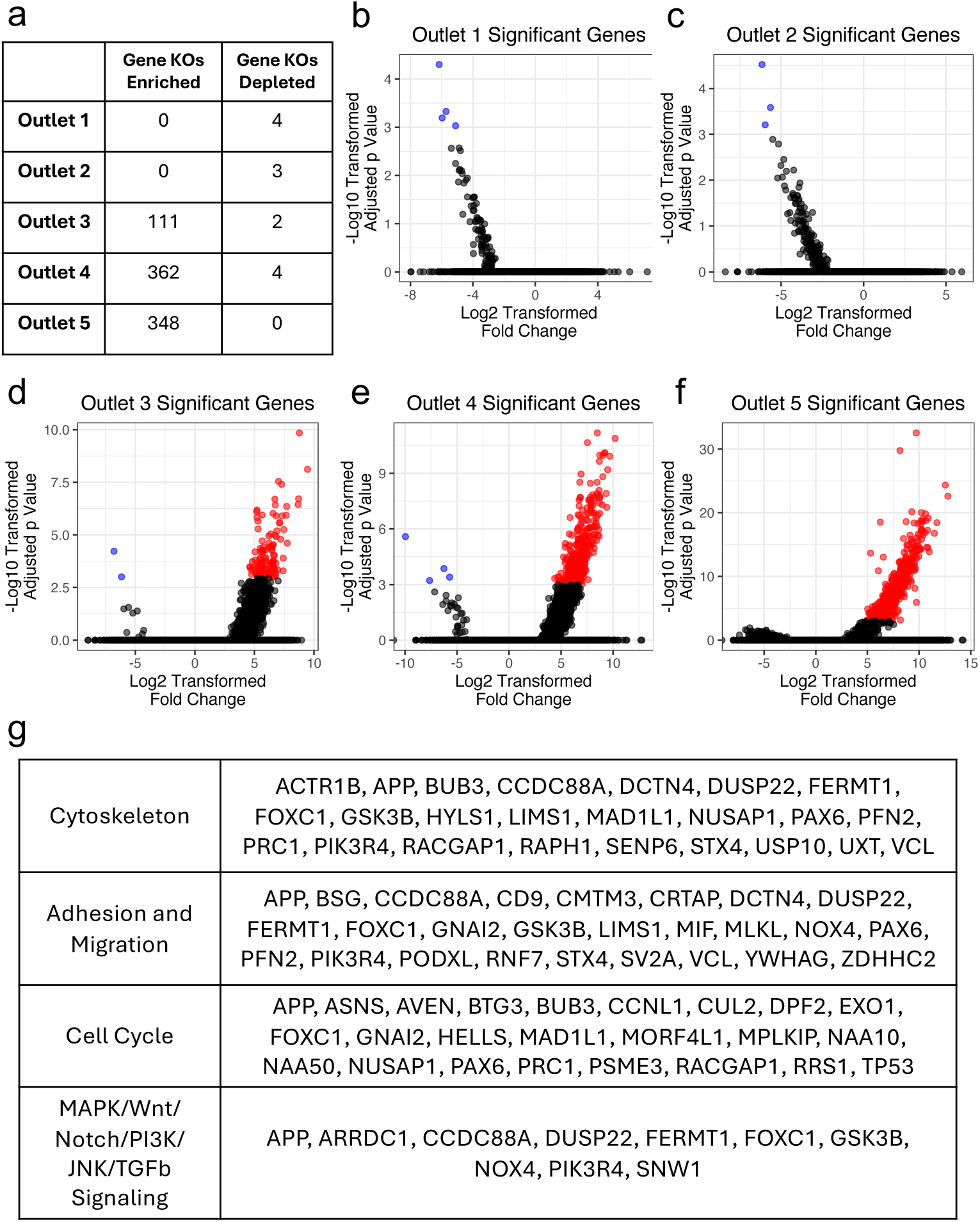
DESeq2 analysis reveals hundreds of genes enriched in mechanically sorted subpopulations. a) Table of significantly enriched/depleted genes according to DESeq2 analysis of sgRNA regions of genomic DNA. Ratio of log-2 transformed fold change and log-10 transformed fold change adjusted p-value for each gene evaluated by DESeq2 differential effect test for b) Outlet 1, c) Outlet 2, d) Outlet 3, e) Outlet 4, and f) Outlet 5 of GeCKO mechanical screen. Each red dot represents a significantly enriched or depleted gene in the outlet compared to the inlet sgRNA distribution (adjusted p-value threshold = 0.001). All black dots represent non-significant genes. g) List of genes with GO terms related to the cytoskeleton, adhesion & migration, the cell cycle, and MAPK/Wnt/Notch/PI3K/JNK/TGFb signaling pathways with a sgRNA fold change over 100-fold and in the top tercile of gene expression in the HEY A8 cell line.

**Supplemental Figure 3.**
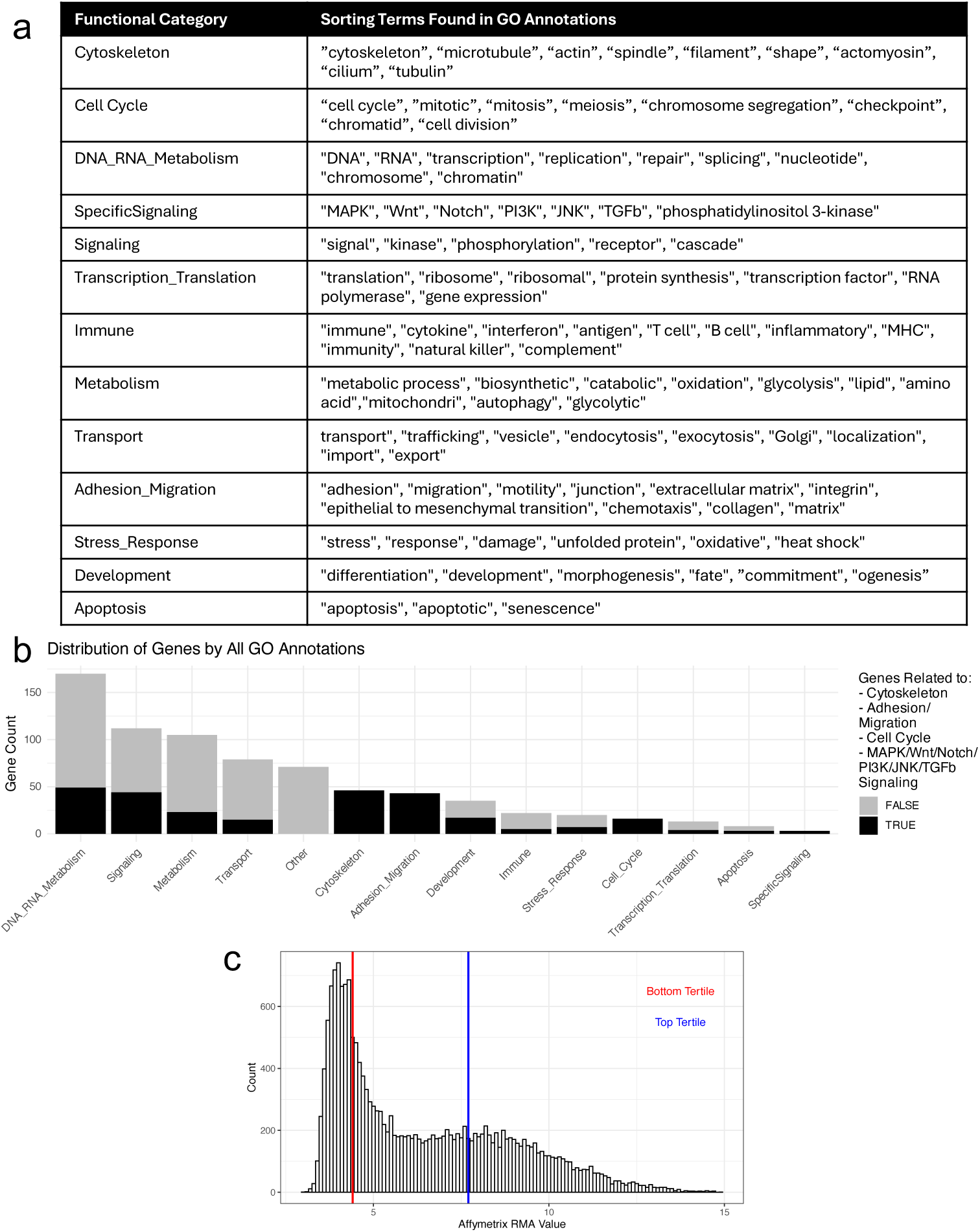
GO annotation mining and population expression shows relevant functional groups and highly expressed genes. a) Table defining functional categories based on presence of sorting terms in GO annotations for biological processes, cellular components, and molecular functions. GO annotations without any of these terms were defined as “Other”. b) Distribution of genes by GO annotations. Main category was defined as the most frequent category for all GO annotations (or second most frequent if the main category was “Other”). Additionally, if any annotations related to the categories of Cytoskeleton, Adhesion & Migration, Cell Cycle, or Specific Signaling were associated with a gene, those genes were flagged. c) Histogram showing genes by expression value in HEY A8 cell line using Affymetrix Gene Chip data from the Gene Expression Omnibus (GSM887080).

### Knockout of PIK3R4, CCDC88A, and GSK3B validated by increased cell stiffness

From the 60 genes identified as associated with the cytoskeleton, adhesion, migration, cell cycle, and specific signaling pathways of interest, that were also highly expressed in HEY A8 cells and whose knockout enrichment in the stiff outlet was greater than 100-fold, we furthermore focused on genes that could be clinically relevant to ovarian cancer patients. Kaplan-Meier (KM) survival analysis for over 600 ovarian cancer patients was conducted to identify genes whose expression is associated with significant changes (p<0.05) in both a patient’s progression-free survival (PFS) and overall survival (OS) from a publicly-available online database^35^. After identifying from a principal components analysis of the sgRNA enrichment analysis that the Outlet 4 samples cluster distinctly from all other samples while the Outlet 5 samples cluster with the Outlet 3 samples (Supplemental Figure 4a), we additionally decided to narrow our genes of interest list to gene knockouts enriched in Outlet 4. Applying these two further filters (significant changes in ovarian cancer patient PFS and OS, and gene knockouts enriched in Outlet 4) reduced our list of genes of interest to 6 candidates – *CUL2, CCDC88A, GSK3B, NAA50, PIK3R4*, and *SV2A* (Figure 4a). Each of these genes when highly expressed (top tercile) in ovarian patients was associated with worse prognosis for patients, with decreases in median PFS of 3.6 – 8.9 months and decreases in median OS of 4.0 – 15.6 months when compared to PFS and OS values of patients who lowly express each of these genes (bottom tercile) (Figure 4a-d, Supplemental Figure 4b-d). Because of their involvement in multiple categories of interest (cytoskeleton, adhesion and migration, and specific signaling pathways), we decided to validate the role of *PI3KR4, CCDC88A*, and *GSK3B* in cell stiffness. After transfecting HEY A8 Cas9-GFP cells with sgRNAs against *PI3KR4, CCDC88A*, and *GSK3B*, we observed significantly increased stiffness of cells for all three genes of interest (Figure 4e).

**Figure 4.**
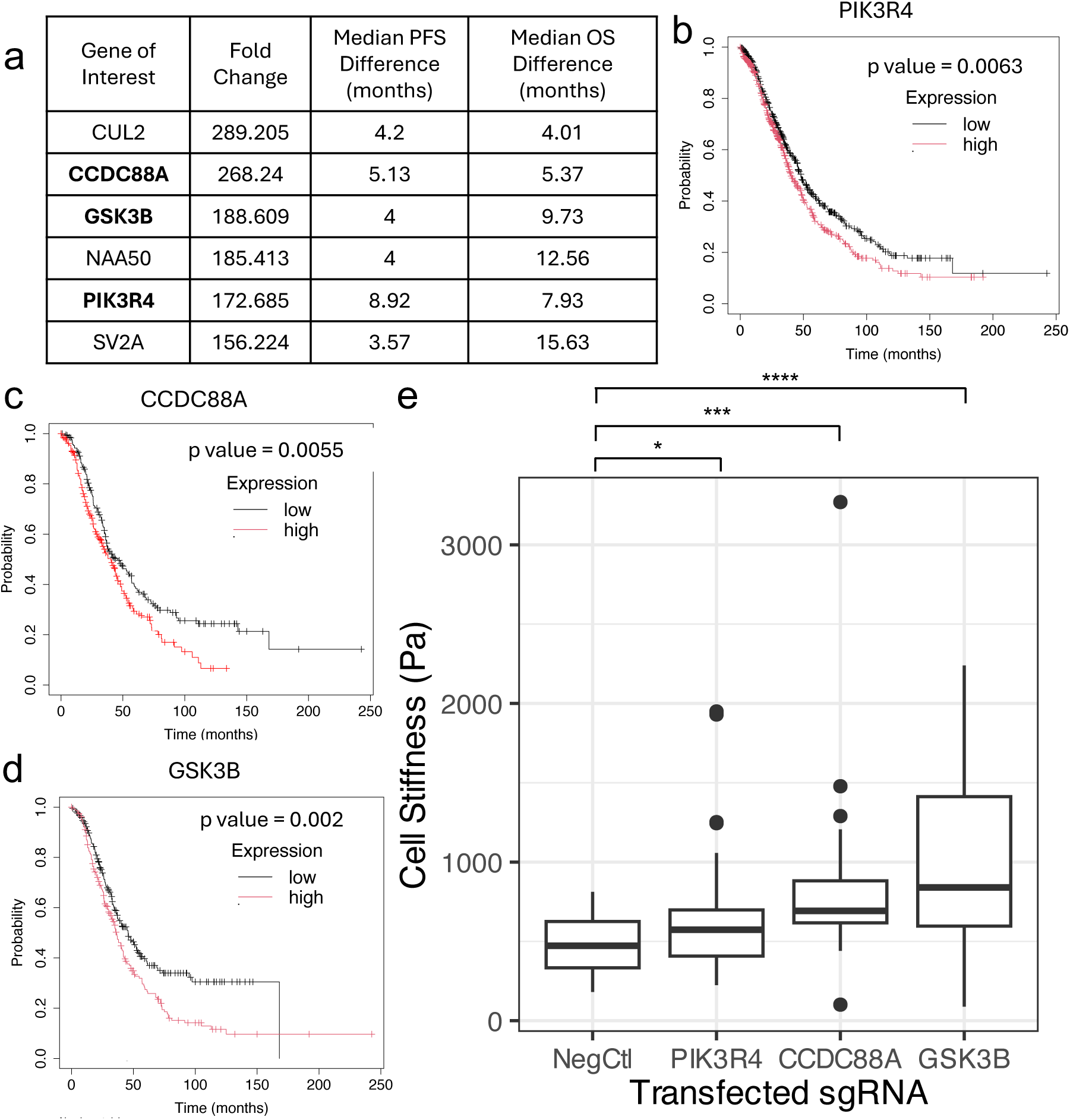
Knockout of PIK3R4, CCDC88A, and GSK3B found to be enriched in stiff outlets and confirmed increased cell stiffness. a) A set of genes of interest that were enriched in Outlet 4 and result in a significant (p < 0.05) difference in both overall survival (OS) and progression free survival (PFS) according to Kaplan-Meier survival analysis of ovarian cancer patients. Table includes the fold change enrichment in outlet 4 and difference in survival in months for OS and PFS (with low expression resulting in better prognosis for all listed genes). Bolded genes were chosen for confirmation experiments. Kaplan-Meier overall survival curves shown for b) *PIK3R4,* c) *CCDC88A,* and d) *GSK3B*. e) AFM data showing knockouts of genes of interest resulted in significant stiffening of cells compared to a negative control. (*, p < 0.05, ***, p < 0.001, ****, p < 0.0001, Welch two-sample t-test, n = 40 per condition)

**Supplemental Figure 4.**
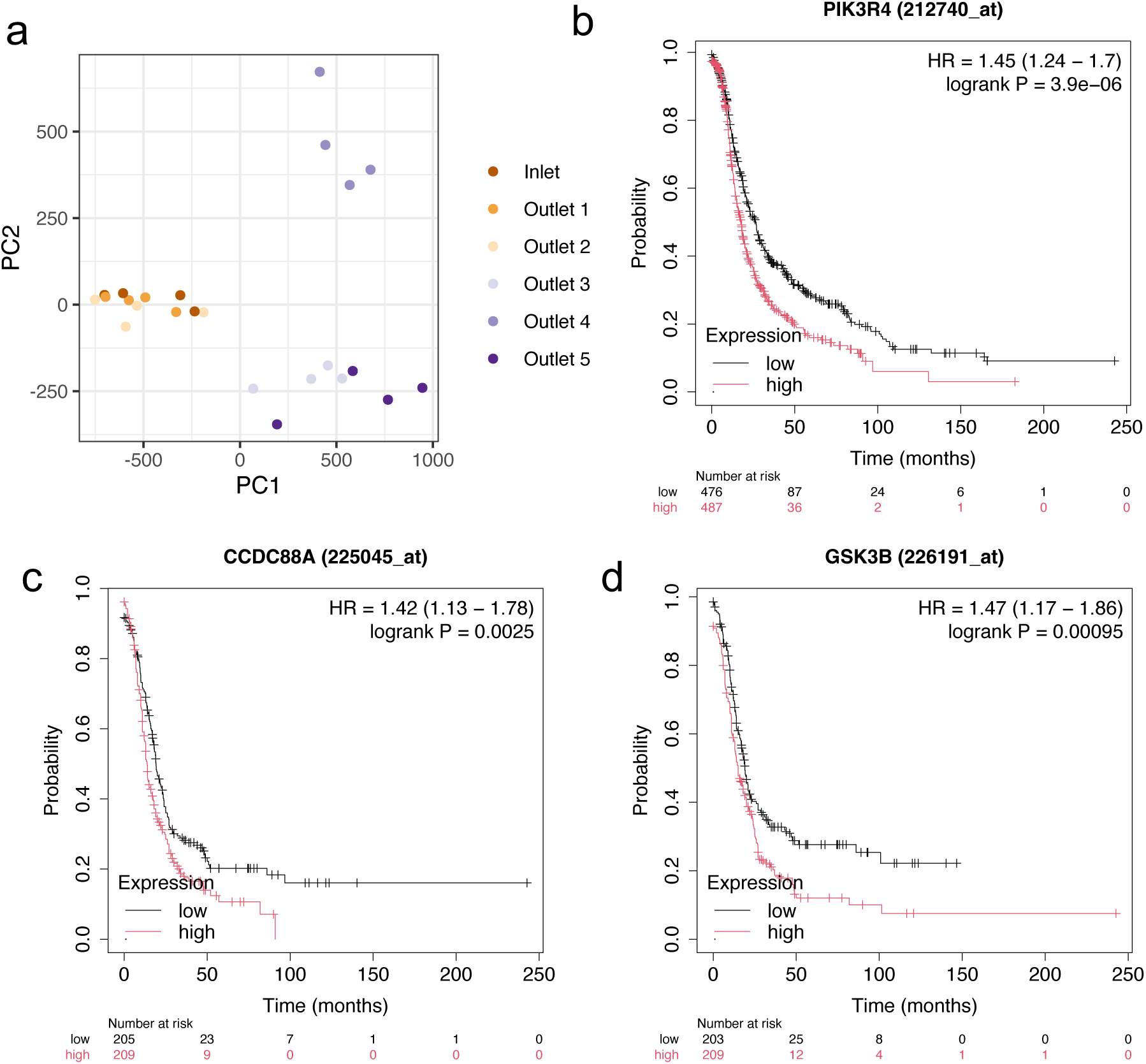
Determining genes of interest for validation. a) Principal component analysis showing clustering of inlet, outlet 1, and outlet 2 samples, clustering of outlet 3 and outlet 5 samples, and distinct clustering of outlet 4 samples. This result motivated the selection of genes enriched in outlet 4 for follow up confirmational experiments. Kaplan-Meier progression free survival curves for b) *PIK3R4,* c) *CCDC88A*, and d) *GSK3B* expression in ovarian cancer patients.

## Discussion

Using a high-throughput, multiplexed microfluidic cell sorting device, we completed the first ever genome-scale CRISPR screen for cell mechanics in human cells. Over 150 million cells possessing single gene knockouts that spanned the entire protein-coding genome were sorted into five mechanical subsets. Analysis of the gene deletions associated with each subset resulted in a large linked dataset correlating gene knockouts to the cell softening and cell stiffening processes. Hundreds of significant genes were found to be enriched or depleted in the five mechanical subsets, providing a large number of hypotheses for future testing of potential genetic control points for cell mechanotype and phenotype. The high throughput genomechanical screening platform was validated by the analysis of the knockdown of *PI3KR4, CCDC88A,* and GSK3B and confirmation of increased stiffness as measured with AFM. This work helps establish cell stiffness as a selectable phenotype analogous to viability, drug resistance, or migration. Furthermore, the identification of potential targets increasing cell stiffness can potentially be used to improve patient prognosis.

Interestingly, we observed that gene knockouts across the genome resulted in an average stiffening of cells (Figure 2b) as well as asymmetrical number of knockouts resulting in cells sorted to the stiff outlets as compared to those sorted to the soft outlets (Figure 3). This possibly could be due to a baseline softness of the highly metastatic HEY A8 cancer cell line which could be near a mechanical minimum value already optimized for migration and invasion^8^. Additionally, the mechanism of sorting used by the microfluidic device design emphasizes enrichment of stiff cells versus softened cells^29^. To further explore gene knockouts that are associated with processes that actively soften cells, further optimization of the flow parameters to adjust the sorting efficiency or alternative sorting methods could be used, such as microbarrier^36^, ratchet^37^, spiral microchannel^38^, real-time deformability cytometry^39^, and deterministic lateral displacement^40^ devices. It is also possible that biologically there is more redundancy in cell softening pathways so that a single gene knockout does not result in active cell softening, in contrast to cell stiffening pathways that are directly linked to disruption of single genes.

While we chose to focus the validation of our method with gene knockouts related to structures and processes conventionally associated with cell mechanics, such as the cytoskeleton, adhesion, migration, cell cycle, and certain signalling pathways, there are a number of other surprising relationships to explore between gene functions and cell mechanics (Supplemental Figure 3b). Notably, a number of gene knockouts related to metabolism were enriched in the stiff outlets. A deeper pursuit of mechano-metabolism is warranted, with particular interest in understanding how cells can tune energy production and internal mechanics in response to external mechanical cues^41–45^. The assays developed here provide a rich area for future study testing hypotheses connecting cell mechanical properties with genes related to metabolism, mitochondria, and energy production beyond the cytoskeletal genes we focused on in this current study, *PI3KR4, CCDC88A*, and *GSK3B*.

*PI3KR4* was one gene knockout identified by the mechanical screen that was confirmed to have a role in cell mechanics. *PI3KR4* encodes for a phosphoinositide-3-kinase regulatory subunit, also known as p150 or VPS15, that specifically directs PI3K activity towards autophagy, endosomal trafficking, and cytokinesis^46,47^. Interestingly, high expression of *PI3KR4* in chronic lymphocytic leukemia has been associated with poor prognosis, representing another cancer type beyond our findings from Kaplan-Meier curve analysis for ovarian cancer patients associated with poor survival statistics^48^. *PI3KR4* mutations have also been associated with perturbations in cell migration and regulation of primary cilium lengths^49,50^. This draws a further connection between this gene of interest, microtubule and cytoskeletal organization, and migratory abilities, further supported by our validation that loss of *PI3KR4* results in cell stiffening.

The second gene knockout we explored that was significantly enriched in the stiffness screen was *CCDC88A,* which encodes for coiled-coil and HOOK domain protein 88A, also known as Girdin, GIV, or APE. Girdin binds to and promotes the crosslinking of actin filaments, plays an important role in migration and lamellipodia formation, and is regulated through phosphorylation by Akt in response to migratory cues and growth factors^51^. Girdin protein expression has been associated with poor prognosis for a number of invasive cancer types, including colon, lung, and breast, primarily because of its role in cell motility^52–54^. Girdin specifically has been linked to cell orientation and directed migration through actin cytoskeleton organization and Golgi body positioning^55^. While it is perhaps surprising that a knockout of a protein known for girding actin filaments would result in global cell stiffening, a potential explanation is that stiffening of lamellipodial region may lead to increased compliance of perinuclear regions where global stiffness measurements were taken. This study supports further exploration of how proteins associated with force generation and cytoskeletal remodeling could also be related to migratory cell softening.

The final gene knockout associated with cell stiffening we studied was *GSK3B* which encodes for glycogen synthase kinase-3 beta, which acts as a negative regulator of glucose homeostasis and Wnt signaling. Of particular interest, *GSK3B* has been associated with the negative regulation and stabilization of microtubules, with reports of ErbB2-repression of GSK3 activity leading to microtubule capture and stabilization particularly at the cell’s leading edge^56,57^. Specifically in cancer, *GSK3B* has been shown to regulate the epithelial-to-mesenchymal transition and poor prognosis for patients with triple negative breast cancer – of note is that GSK3B inhibitors selectively killed mesenchymal, stem-like cancer cells^58^. Our validation of cancer cell stiffening after *GSK3B* knockout which is associated with microtubule stabilization and prevention of an epithelial-to-mesenchymal transition is in accordance with a hypothesis linking a softer mechanotype with more mesenchymal and more migratory phenotypes, and a potential target for control of cell behavior.

There are limitations to the GeCKO mechanics method described here that could be further explored and amended in the future. Using a low MOI for the lentiviral transduction results in most cells receiving at most one gene knockout. However, the number of cells receiving two or more integrants could be close to 20% of all infected cells when using an optimized infection efficiency of 40%^59^. With the current sequencing protocol, we are unable to determine whether the sgRNA that ends up in each outlet was the only integrant in a cell or a multi-integrant. However, because of the redundancy of using ∼500 cells per sgRNA, 4-6 sgRNAs per gene, and 4 bio-replicates, we can say with more confidence that the genes that appear significantly enriched in an outlet are most likely due to the knockout of that specific gene and not a combination of genes. Additionally, the GeCKO mechanics method should be viewed primarily as a hypothesis generation tool and all potential genes correlated with each mechanical subset need to be validated. It should also be noted that we chose to demonstrate this method with a single ovarian cancer cell line though there is great potential for expanding this analysis to additional ovarian cancer cell lines, other cancer types, and other cell types to identify more broad genetic controllers of cell mechanics. Lastly, because of the design of the microfluidic sorting platform, mechanical phenotype is distinguished under specific flow and deformation conditions, requiring cells to be sorted while in suspension, and cannot distinguish between the cortical, nuclear, and cytoskeletal contributions to global cell stiffness. This method may be particularly applicable to studying the genomechanical relationships of cells that are found in suspension in the body, such as blood cells or metastasizing cells.

Finally, there are a number of interesting and exciting directions this work could take next. This same method could be used to mechanically screen genes of interest in other cell types as a comparison between cancer types with distinct metastatic potential or between cancer cells and stromal cells. Exploring these questions could accelerate the finding of sets of universal genes that are associated with the mechanical properties of all cells. Additionally, there are a number of other GeCKO libraries that have already been developed that could also be substituted for the Brunello library we used in this study. For example, there are libraries of sgRNAs to knock out epigenetic regulators and lncRNAs as well as libraries that activate, instead of knock out, genes of interest. This procedure could be repeated with different combinations of cell types and sgRNA libraries of interest to generate more testable hypotheses about the connection between cell phenotype, genotype, and mechanotype. Overall, the enabling technology we describe can expand the CRISPR screening paradigm from molecular phenotypes to biophysical markers to identify potential genetic regulators of mechanotype and phenotype.

## Methods

### Development of CasG-expressing cell line

We have developed an ovarian cancer cell line that expresses Cas9 allowing for delivery of sgRNA to knock out particular genes of interest. HEK293T cells (provided by Lena Gamboa in Dr. Gabe Kwong’s lab) were cultured on poly-L-lysine (Sigma Aldrich) coated petri dishes (VWR) and transfected at 80-95% confluence with a lentiviral transfer vector containing Cas9 from *Streptococcus pyogenes* and GFP for detection of vector integration, psPAX2, VSV-G, TransIT-LT1transfection reagent (Mirus Bio), and Opti-MEM (ThermoFisher). Three days after transfection, viral particles were harvested and concentrated using overnight incubation at 4 ℃ with PEG-it (System Biosciences) and centrifugation at 1500xg for 30 minutes. Viral particles were resuspended in PBS and stored at −80℃ until use. Ovarian cancer cell HEY A8 cells (provided by Dr. John McDonald) were transduced with the varying amounts of the concentrated lentiviral particles (0, 10, 20, 40, and 60 µL per confluent 24-well), removing the virus the following day. Each well was trypsinized and resuspended for single cell sorting with FACS (BD FACS Aria) into a 96 well plate. Cells transduced with 0 µL of virus were used to create a GFP-negative gate. We additionally applied a high-GFP and mid-GFP gate, selecting the mid-GFP gate for the single cell sort, because we wanted to select cells that had GFP (and thus Cas9) integrated into their genome, but not too frequently (because of unwanted issues with the locations the vector integrates into the genome). Single cells deposited in 96-well plates were expanded into clonal populations over two weeks.

To test the efficacy of the expressed Cas9, we transfected the HEY A8-Cas9-GFP cells with 1 µg/µL sgRNA against GFP (sgGFP sequence: GGGCGAGGAGCTGTTCACCG) with Lipofectamine 2000 (ThermoFisher) in OptiMEM (ThermoFisher) for 24 hours. Five days after transfection, flow cytometry (BD Accuri) was used to compare the FL1 expression of HEY A8-Cas9-GFP cells with and without the addition of sgGFP. Additionally, DNA from the transfected cells was extracted using Quick Extract (Lucigen) and the GFP region was amplified with PCR. Gel electrophoresis was used to confirm an amplified GFP region and PCR products were sent for Sanger sequencing (Eton) to determine the insertion-deletion rate after Cas9 editing using Tracking of Indels by Decomposition (TIDE) analysis (https://tide.nki.nl).

### Cell culture and sample preparation

The HEY A8-Cas9-GFP ovarian cancer cell line was cultured in complete media consisting of RPMI-1640 (Sigma-Aldrich) supplemented with 10% v/v fetal bovine serum (Atlanta Biologicals) and 1% v/v penicillin-streptomycin (Sigma-Aldrich). HEK293T cells were cultured in complete media consisting of DMEM (ThermoFisher) with 10% v/v fetal bovine serum (Atlanta Biologicals) and 1% v/v penicillin-streptomycin (Sigma-Aldrich). Cells were cultured at 37℃ at 5% CO_2_.

For the GeCKO mechanics microfluidic sort, cells were prepared for processing by trypsinization and resuspension in flow buffer (complete media, 10% v/v OptiPrep (Sigma-Aldrich) and 0.1% v/v Pluronic F-127 (Millipore Sigma)) with 1 mg/mL DNAse I (Sigma-Aldrich) at approximately 1-2 million cells/mL. The formulation of OptiPrep in the flow buffer was optimized to create a neutrally buoyant solution to prevent cell settling in the device or tubing. In short, cells were resuspended in solutions with different concentrations of each density gradient medium and centrifuged at 300xg for 10 minutes and the concentration at which a cell pellet first began to form was selected as the ideal buffer density. Cells were strained using a 20 µm pore filter (PluriSelect) and loaded into a syringe (VWR) for microfluidic processing.

### GeCKO viral library transduction

The human CRISPR Brunello lentiviral pooled library was designed to optimize on-target activity and reduce off-target effects (Addgene 73178-LV)^25^. This library targets 19,114 human genes with 76,441 total gRNAs, with 3-4 gRNAs targeting each gene. The 2-vector system was selected where the Cas9 component and guide library are delivered in different viral vectors. This allows for the reduction of noise in the knockout screen, a higher-titer guide lentivirus for a more efficient transduction, and a recombinant system that is safer for use in the lab. To maintain coverage of approximately 315 cells with each sgRNA knockout after transduction, approximately 60 million cells were transduced with the lentiviral library. Cells were seeded at a high concentration of 3 million cells per 12-well in RMPI supplemented with 10% v/v heat-inactivated FBS, 1 µg/µL polybrene (Sigma Aldrich) and 300 µL of the lentiviral library (2.88E6 TU/mL). Transducing the Cas9-expressing cells with a low MOI (0.3-0.5) ensures that statistically most cells will only receive at most one knockout^59^. The GeCKO infection protocol was optimized by transducing the HEY A8-Cas9-GFP cells with varying amounts of the lentiviral library (0, 50, 100, 300, and 500 µL). Puromycin was used as selection agent to isolate the cells that successfully received a single gene knockout. We used 0.5 µg/mL puromycin as the minimal antibiotic dose that killed all cells after 1 week of treatment. After 1 week of puromycin selection, 40% of the cells that had been transduced with 300 µL of lentiviral library survived when compared with cells that did not undergo puromycin selection, the optimal infection efficiency. For the GeCKO sort, the cells that were transduced with the lentiviral library and underwent puromycin selection were expanded to provide at least 40 million cells per microfluidic processing to maintain coverage of all sgRNAs.

### AFM and force curve analysis

To investigate the effect of transduction of the GeCKO library on cell mechanics, we used force spectroscopy to obtain force-indentation curves with an atomic force microscope (Asylum Research) with an integrated optical microscope (Nikon) on a vibration isolation table. For improved contact area between the cantilever and cell, a 7.32 μm spherical polystyrene particle was attached to a tipless silica nitride cantilever (Bruker Probes) using a two-part epoxy (JB Weld). Before taking measurements, the cantilever was calibrated to determine the deflection inverse optical lever sensitivity and spring constant (k was approximately 30 pN/nm).

Cells were seeded after trypsinization in a glass bottom FluoroDish (World Precision Instruments) and allowed to attach before force curve collection. For measurements, the cantilever probe was visually aligned with the cell center and moved with a velocity of 2 μm/s to indent the cell with increasing compressive force until a force trigger of 10 nN was reached. The cantilever was held in position for 5 seconds, dwelling towards the surface, allowing for viscous relaxation of the cell before reversing the direction of its velocity. We collected 150 cellular measurements per cell type.

We used custom code written in R (https://github.com/nstone8/Rasylum) relying on the Hertzian contact model, which describes non-adhesive elastic contact between two bodies, to calculate the cellular reduced Young’s modulus^60^. The contact point was estimated by the intersection of the flat, undeformed region of the force curve with a line fit to the force curve region where the cantilever was in contact with the cell. Next, we identified the true contact point by iteratively testing the points around the estimated contact point with the minimal residual difference between the measured force curve and a nonlinear fit described by the governing equation Hertz used to describe the contact between an elastic sphere and an elastic half space.

### High-speed video analysis

High-speed videos were taken of the cells passing through a single channel sorting device operating at 30 µL/min total flow rate using a Phantom camera at 2500-3000 frames per second to monitor cell trajectories and velocity after interactions with the diagonal ridges. To measure cell trajectories, ImageJ was used to extract an image stack from each video and a Z-projection was used to identify the background of each image. The calculator plus function was used to subtract the background from the image stack so that only moving cells would remain. Using the TrackMate^61,62^ plugin in ImageJ^63,64^, cells were identified in each frame and linked together into tracks tracing each cell’s movement through the device. This produced approximately 100 tracks per video. Custom python software (https://github.com/nstone8/heimdall) was used to analyze collected cell trajectories. The tracks of transduced and non-transduced cell populations were analyzed to determine deflection and interaction time with each ridge.

### Microfluidic mechanical screen

The library of single gene knockout cells was processed through the multiplexed microfluidic cell mechanical sorting device described in chapter 4. The microfluidic device was set up on the stage of an optical microscope (Nikon) to monitor cell sorting. First, the device was primed by flowing flow buffer through the device to remove all air bubbles. Flow was controlled using two syringe pumps (Harvard Apparatus) with 0.02 in inner diameter and 0.06 in outer diameter Tygon tubing (Cole Parmer) connecting the sheath flow syringes of flow buffer to the inlet ports of the left and right sheath channels. The two independent sheath flows allow for the control of cell focusing at the entrance of the sorting channel. The tubing fits directly into the 1.2 mm punch without need for adapters. All tubing and needles used in the experiment were cleaned prior to use with deionized water. Tubing was also fit into the five outlet ports and placed in a conical tube for sorted cell collection. Once the device was prepared, the syringe of cells in flow buffer with DNAse were connected to the cell inlet of the device with tubing using a third syringe pump. The flow parameters were optimized (see 4.3.7) for sorting of the ovarian cancer cells based on stiffness, with 50 µL/min total flow rate per channel, or 200 µL/min and 120 µL/min sheath flows, and 80 µL/min cell inlet flow for the entire multiplex device. The optimal device gap size was 9.5 µm with 14 diagonal ridges. After distribution, cell sorting, and recombination of fluid in the multiplexed device, cells were separated into five subsets – softest, softer, median, stiffer, and stiffest cells. Cells were collected in conical tubes and kept on ice until the sort was complete. After recording the volume and concentration of cells collected at each outlet, cells were pelleted and flash frozen at −80℃ for genomic DNA extraction. The GeCKO mechanical screen was repeated 4 times with 4 biological replicates.

### Isolation and purification of genomic DNA

Genomic DNA was isolated and purified using the Wizard Genomic DNA Purification Kit (Promega). In short, pelleted cells were washed in PBS and lysed using a nuclei lysis solution with RNAse. After incubation for 15 minutes at 37℃, a protein precipitation solution was added and DNA from the supernatant layer was transferred to a clean tube. The addition of isopropanol (Sigma Aldrich) precipitated the DNA which was washed with 70% ethanol (Fisher Scientific), pelleted, and air dried before resuspension in a DNA rehydration solution. DNA was stored at 4℃ and a Nanodrop spectrophotometer (ThermoFisher) was used to determine the quantity and quality of DNA from each sample.

### Genomic DNA sequencing

Next generation sequencing (NGS) was used to confirm the sgRNA distribution after the screen. First, we prepared our library for sequencing using PCR to amplify the sgRNA target region (P5 primer: ACACTCTTTCCCTACACGACG CTCTTCCGATCTTTGTGGAAAGGACGAAACACC*G, P7 primer: GACTGGAGTTCAGACGTGTGCTCTTCCGATCTTCTACTATTCTTTCCCCTGCACTG*T, product size: 354 nucleotides) of gDNA from each sample (five outlets and one inlet for each of the four bio-replicates) with a second PCR step to add Illumina adaptor sequences and barcodes for batch NGS. Next, the PCR products were purified and quantified using a Bioanalyzer (Agilent) before combining and sequencing the indexed samples for sgRNA distribution on a NovaSeq 6000 (Illumina) using the SP 100 cycle kit with 122 cycles of read 1 (forward) and 8 cycles of index 1 with a coverage of at least 100 reads per sgRNA in the library (650-800 million reads total).

### Screen analysis

The web-based Basespace Sequencing Hub (Illumina) was used to generate FASTQ files from the raw NovaSeq data which were then input into a fully-interactive analysis and documentation suite for pooled CRISPR screens, CRISPRAnalyzeR (https://github.com/boutroslab/CRISPRAnalyzeR). This web-based, interactive analysis suite took the input of the NGS raw data and a description of the Brunello pooled CRISPR sgRNA library and output the data quality, read count distribution, replicate quality and sample comparison, and sgRNA statistics. After passing this screen quality step, the software completed six differential analysis methods (Wilcox, DESeq2, MAGeCK, sgRSEA, EdgeR, and Z Ratio) and two Bayesian analysis methods (BAGEL and ScreenBEAM) comparing sgRNA distributions of each outlet to the inlet distribution. We focused our analysis on the DESeq2 method which uses the summed read counts of all 4-6 sgRNAs per gene and tests for differential effects between the inlet and outlet distributions using a negative binomial distribution model^65^. The total read count per gene is normalized, size-factors are estimated, and variance is stabilized with a parametric test. A Wald test is used to compare the difference in fold changes in the inlet and outlet distributions to determine whether the gene was enriched or depleted. We used an adjusted p-value threshold of 0.001 to determine significance for the DESeq2 method.

### Quality control of sequencing analysis

For quality control of the sequencing data, FastQC was used to determine the mean quality value across each base position in the read after removing the Illumina Adapter sequence (TTGTGGAAAGGACGAAACANNN). MAGeCK count was used to determine the Gini Index for each sample while MAGeCK test was used to generate the heatmap, hierarchal clustering, and principal components analysis of each sample^66,67^.

### Identifying genes of interest

Gene ontology annotation was performed using the functional categories defined in Supplemental Figure 3a to sort GO annotations across biological processes, cellular components, and molecular functions. In particular, genes related to the cytoskeleton, adhesion & migration, cell cycle, or the MAPK, Wnt, Notch, PI3K, JNK, and TGFb signaling pathways were flagged. Total gene expression in the HEY A8 cell line was determined using publicly available Affymetrix Gene Chip data from the Gene Expression Omnibus (GSM887080) and expression was divided into top, middle, and bottom terciles. Kaplan-Meier analysis was completed using a web-based plotter (kmplot.com) using mRNA gene chip data for ovarian cancer patients^35^. Patients were split by trichotomization comparing the lower and upper tercile of expression and only JetSet best probe sets were used^68^. Analysis was not restricted by histology, stage, grade, or TP53 mutation. After determining genes of interest for validation, the following sgRNAs were used to knockout each gene of interest: *PIK3R4* – CAAGAACCAGATGACAAACG, *CCDC88A* – GGTTGCCGCACATATTCAAG, and *GSK3B* – GTGGCTCCAAAGATCAACTC.

## Data and Code Availability

AFM force curves and the CRISPR screen sgRNA sequencing will be shared by the corresponding authors upon request. Custom code used for AFM force curve analysis, high speed video cell tracking, and CRISPR screen analysis can be found at https://github.com/nstone8/Rasylum, https://github.com/nstone8/heimdall, https://github.com/boutroslab/CRISPRAnalyzeR. Any additional information required to reanalyze the data reported in this paper is available from the corresponding authors upon request.

## Acknowledgements

This research project was supported by through the National Institutes of Health: National Cancer Institute, grant #F31CA243345, and National Institutes of Health: National Cancer Institute, grant #F32CA281162. We wish to acknowledge the core facilities at the Parker H. Petit Institute for Bioengineering and Biosciences at the Georgia Institute of Technology for the use of their shared equipment, services, and expertise, specifically the High Throughput DNA Sequencing Core and Cell Analysis and Cytometry Core.

## Author Contributions

**Katherine M Young:** conceptualization; data curation; formal analysis; funding acquisition; investigation; methodology; software; validation; visualization; writing – original draft preparation; writing – review and editing, **Curtis Dobrowolski:** data curation; formal analysis; software; visualization, **Nicholas Stone:** data curation; formal analysis; investigation; software; visualization, **Kalina Paunovska:** formal analysis; investigation; visualization, **Sydney Bules:** investigation, **Kelly Ahkee**: investigation, **James Hankish:** investigation, **Alex Chapman:** investigation, **James Dahlman:** conceptualization; resources; supervision, **Todd A Sulchek:** conceptualization; funding acquisition; project administration; resources; supervision; writing – review and editing, **Cynthia A Reinhart-King:** conceptualization; funding acquisition; project administration; resources; supervision; writing – review and editing

